# Evaluation of K18-*hACE2* mice as a model of SARS-CoV-2 infection

**DOI:** 10.1101/2020.06.26.171033

**Authors:** G. Brett Moreau, Stacey L. Burgess, Jeffrey M. Sturek, Alexandra N. Donlan, William A. Petri, Barbara J. Mann

## Abstract

Murine models of SARS-CoV-2 infection are critical for elucidating the biological pathways underlying COVID-19 disease. Because human ACE2 is the receptor for SARS-CoV-2, mice expressing the human *ACE2* gene have shown promise as a potential model for COVID-19. Five mice from the transgenic mouse strain K18-*hACE2* were intranasally inoculated with SARS-CoV-2 Hong Kong/VM20001061/2020. Mice were followed twice daily for five days and scored for weight loss and clinical symptoms. Infected mice did not exhibit any signs of infection until day four, when weight loss, but no other obvious clinical symptoms were observed. By day five all infected mice had lost around 10% of their original body weight, but exhibited variable clinical symptoms. All infected mice showed high viral titers in the lungs as well as altered lung histology associated with proteinaceous debris in the alveolar space, interstitial inflammatory cell infiltration and alveolar septal thickening. Overall, these results show that symptomatic SARS-CoV-2 infection can be established in the K18-*hACE2* transgenic background and should be a useful mouse model for COVID-19 disease.

## Introduction

An invaluable step in identifying effective vaccines and therapies to combat COVID-19 is the availability of a mouse model of infection. The host receptor for SARS-CoV-2 is the human angiotensin-converting enzyme 2 (hACE2) (Wan *et al*., 2020), which was previously identified as the receptor for the SARS-CoV-1 virus (Li *et al*., 2003) that causes Severe Acute Respiratory Syndrome (SARS), a disease that emerged from China in 2002 (WHO, Tsang). The mouse ACE2 ortholog, which has significant amino acid sequence variation in the viral receptor binding domain, cannot serve as an efficient receptor for either SARS-CoV-2 or CoV-1 (Shang *et al*., 2020). To develop a mouse model to study SARS-CoV-1 infection, McCray *et al*. developed a transgenic mouse line, K18-*hACE2*, which expresses the *hACE2* gene under the control of the human cytokeratin 18 promoter. Infection of these mice with SARS-CoV-1 results in a rapidly lethal infection that spreads from the lungs to the brain, and induces proinflammatory cytokines and chemokines (McCray *et al*., 2007). Four other *hACE2*-expressing mouse lines have been created to date and tested for the ability to support SARS-CoV-2 infection (Lutz, 2020). Two lines express the *hACE2* gene under the control of the mouse *ACE2* promotor (Bao *et al*., 2020); one was made using CRISPR/Cas9 technology (Sun *et al*., 2020). The third strain uses the lung ciliated epithelial cell *HFH4* promoter (Menachery *et al*., 2016) (Dinnon *et al*., 2020). An additional approach was to transfect wild type mice with an adenovirus carrying the *hACE2* gene (Hassan *et al*., 2020). Overall, with the exception of the HFH4 mice, in which there was some lethality, infection of these three mouse strains with SARS-CoV-2 results in mild clinical symptoms, and no lethality. Here we report the infection of K18-*hACE2* with SARS-CoV-2. While this infection resembled that of other strains, we observed variable clinical presentation, with some mice exhibiting more severe symptoms than reported using other models. Overall, this work supports the usefulness of K18-*hACE2* transgenic mice as a model for human COVID-19 infections.

## Results and Discussion

To investigate the potential of this transgenic mouse strain as a model for COVID-19 infection, five K18-*hACE2* mice were intranasally inoculated with 8 × 10^4^ TCID_50_ of SARS-CoV-2, while five mice were mock-infected with sterile DMEM. Mice were followed twice daily for five days, and scored for clinical symptoms (weight loss, eye closure, appearance of fur and posture, and respiration). The mock-infected mice did not exhibit any clinical symptoms or experience any weight loss throughout the experiment. Infected mice did not exhibit any measurable clinical symptoms until day four, and these were limited to weight loss. On day five all of the infected mice had lost around 10% of their original weight (Figure 1A) and exhibited variability in other clinical signs of infection, with clinical scores ranging from 3-9 (maximal score 14) (Figure 1B). While two of the infected K18-*hACE2* mice showed only mild symptoms at day five (weight loss, and reduced activity), two mice exhibited piloerection. The most severe mouse had increased respiration, lethargy, and slight eye closure, and met our criteria for euthanasia. Because the study was ended on day five, it is unclear whether the remaining four mice would have recovered if the study was carried past day five.

**Figure 1.**
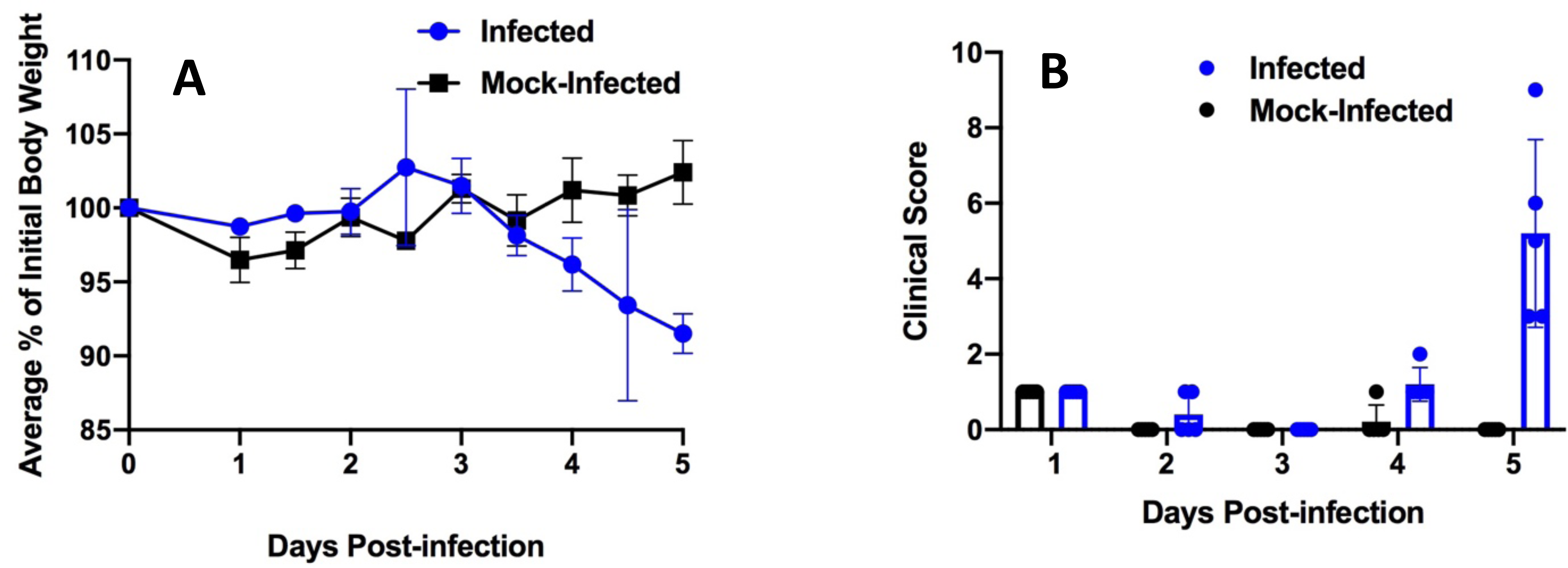
K18*-hACE2* C57Bl/6J mice were intranasally inoculated with 8×10^4^ TCID_50_ of SARS-CoV-2 and weight loss and clinical score were monitored. (A) Weight loss is measured as the percent weight loss compared to the initial weight on day 0. (B) Clinical score consists of weight loss, activity level, eye closure, appearance of fur and posture, and respiration. The mock-infected mice did not exhibit any clinical symptoms or experience any weight loss throughout the experiment.

While the clinical severity was variable between infected K18-*hACE2* mice, our results suggest that these mice present with more symptomatic disease than other *hACE2* mouse models of SARS-CoV-2 infection. In the mouse model expressing *hACE2* under the mouse *ACE2* promoter, infected mice did not exhibit any clinical symptoms other than maximal weight loss on day three post-infection, and those mice recovered (Sun et al, 2020). Only mild ruffling of fur and up to 8% weight loss on day five were observed in the other model using the mouse *ACE2* promoter, and once again all mice recovered (Bao et al, 2020). In mice transfected with an adenovirus carrying the *hACE2* gene mice exhibited about a 10% weight loss on day four post-infection, but no lethality (Hassan *et al*., 2020). In contrast to these models, in which mice exhibited mild symptoms and recovered, only 60% of the mice survived past day five in the mouse strain expressing *hACE2* under the lung ciliated epithelial cell *HFH4* promoter (Dinnon *et al*., 2020). While this model had higher lethality, weight loss was only about 5% and these mice had no respiratory symptoms. The authors hypothesize that mortality may be due to neuroinvasion, as virus was detected in the brain. In K18-*hACE2* mice infected with SARS-CoV-1 the course of infection is clearly different; the infection is uniformly fatal, beginning on day four post-infection, and mice were symptomatic with labored breathing and lethargy. (McCray *et al*., 2007). While the numbers of mice used in this study are small and we were not able to measure survival, our data support a difference in the disease progression between these two viruses.

All mice were euthanized on day five and tissue was collected for dissection and enumeration of viral loads. No significant differences in histology of the spleen, small intestine, or liver were observed between infected and mock-infected mice, and these tissues were normal in size and appearance. Dissection of the lungs of infected mice revealed a mottled or marbled appearance that was not observed in mock-infected mice (data not shown). Lung sections were analyzed after staining with hematoxylin and eosin (H&E) and scored based on tissue pathology (Matute-Bello *et al*., 2011). SARS-CoV-2-infected mice exhibited significantly higher histopathology scores than mock-infected mice (Figure. 2). The major histopathology findings in infected mice were proteinaceous debris in the alveolar space, neutrophils in the interstitial space, and alveolar septal thickening (Figure 2); these observations were consistent with other h*ACE2* mouse models, which also detected signs of lung injury including interstitial pneumonia, inflammatory cell infiltrates, and alveolar septal thickening (Sun et al, 2020; Bao et al, 2020) (Dinnon *et al*., 2020) (Hassan *et al*., 2020). Consistent with the observed infiltrating neutrophils, granulocytes and inflammatory monocytes were also elevated in the bronchoalveolar lavage (BAL) fluid from infected mice (Figure 3).

**Figure 2.**
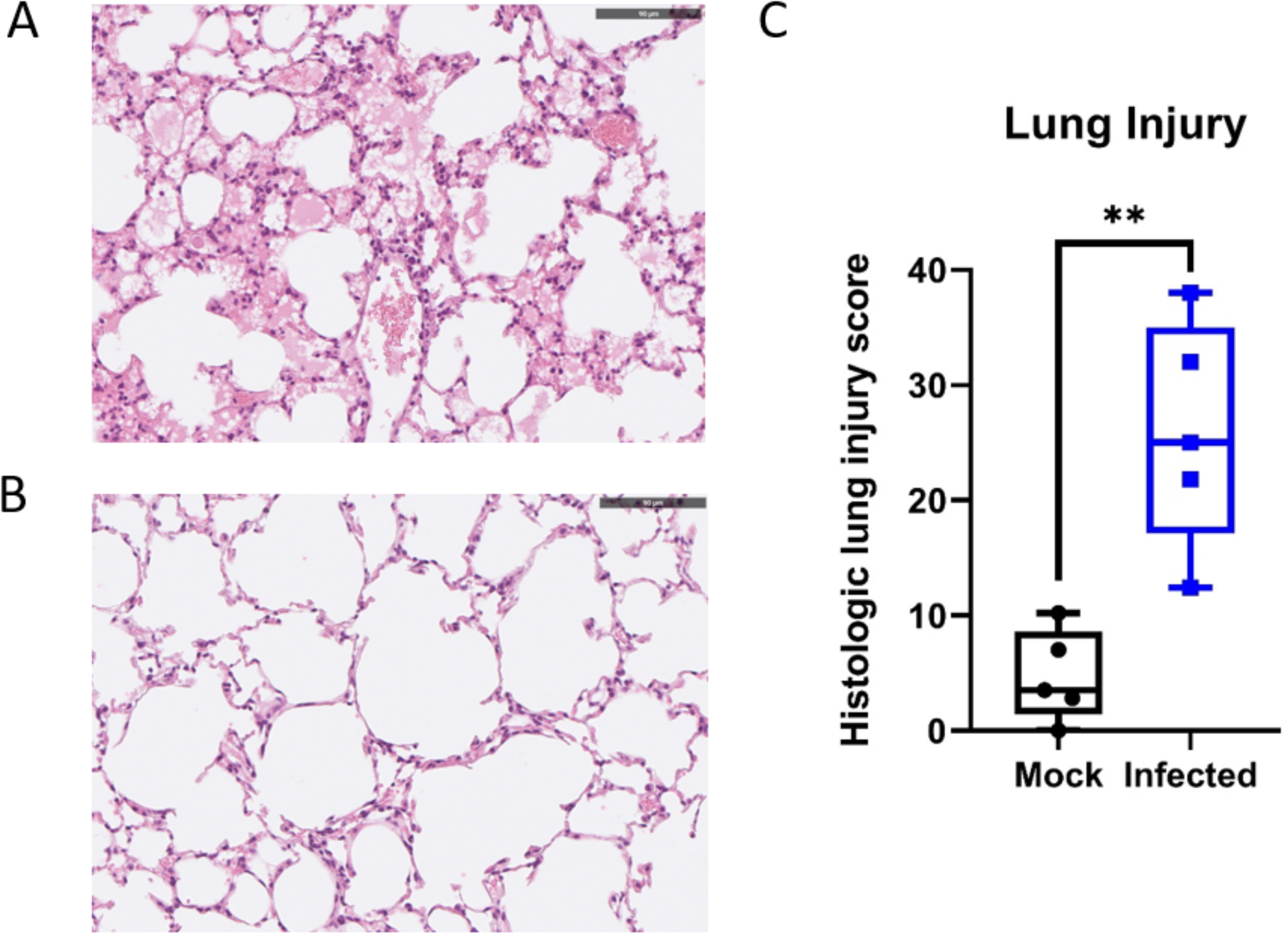
Lung histology of mice infected with SARS-CoV-2. Representative images from H&E stains of from lungs of infected mice (A) and mock-infected (B). Lungs from infected mice had alveolar proteinaceous debris, interstitial inflammatory cell infiltration, and alveolar septal thickening. Blinded quantification of lung injury is shown in C. Scale bar = 90 µm. **p<0.01.

**Figure 3.**
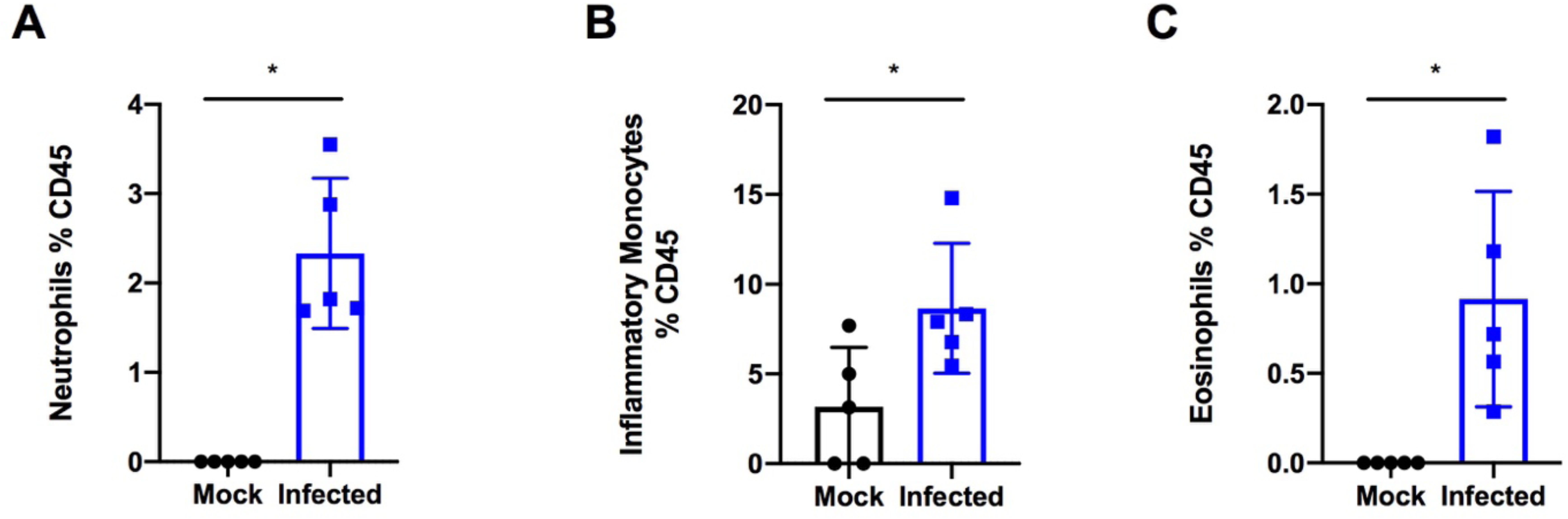
SARS-COV-2 infection increases granulocytes and inflammatory monocytes in bronchoalveolar lavage (BAL) fluid. K18-hACE2 C57Bl/6J mice were intranasally inoculated with 8×104 TCID50 of SARSCoV-2 Hong Kong/VM20001061/2020 (Source: BEI Resources). BAL was collected, cells isolated and stained via flow cytometry. *p<0.05

Other *hACE2* mouse models of COVID-19 infection have observed high viral titers in the lungs with limited viral load in organs such as the liver and spleen during intranasal infection (Sun et al, 2020; Bao et al, 2020). While we did not investigate viral load in the liver or spleen, these organs appeared normal by histology, suggesting that there was limited viral titer in these tissues. Virus was detected in the lungs of all infected mice, with titers generally in the range of 1 × 10^5^ PFU/ml (Table 1). Viral titers in the lungs appeared somewhat associated with disease severity: mouse 1390, which had the highest lung titer, had the highest clinical score, histopathology score, percent weight loss at day five (Table 1), and the highest numbers of neutrophils, monocytes and eosinophils in the BAL (Figure 3). In addition, mouse 1413, which had the lowest titer, had the lowest clinical score, second lowest percent weight loss at day five (Table 1), and lowest number of eosinophils and monocytes in BAL (Figure 3). Of note, mouse 1413 did not have the lowest histopathology score (Table 1). While there were trends towards higher viral titers in the lungs being associated with higher clinical and histopathology scores, these trends were not significant, and viral titer was not a strong predicter of percent weigh loss. The power of this analysis is limited by the small sample size, but these results suggest that factors in addition to viral load, such as inflammatory responses, are driving the severity of disease. This would also potentially explain the sudden onset of clinical symptoms at five days post-inoculation.

**Table 1.** Characteristics of Infected Mice on Day 5 Post-infection.

| Mouse ID | % of Initial Weight (Day 5) | Clinical Score | Average Lung Pathology Score | Viral Load in Lungs (PFU/ml) |
| --- | --- | --- | --- | --- |
| 1413 | 91.6 | 3 | 24 | $7.5 \times 10^3$ |
| 1378 | 91.2 | 3 | 32 | $1.2 \times 10^5$ |
| 1390 | 89.7 | 9 | 38 | $1.75 \times 10^5$ |
| 1387 | 91.6 | 5 | 12.4 | $5.0 \times 10^4$ |
| 1382 | 93.5 | 6 | 21.8 | $1.5 \times 10^5$ |

In this report we have described the course of SARS-CoV-2 infection in K18-*hACE2* transgenic mice. Our findings are consistent with other studies utilizing *hACE2* mice, which observed successful infection with SARS-CoV-2 and a milder disease severity compared to SARS-CoV-1 (Bao *et al*., 2020)(Sun *et al*., 2020). The onset of symptoms was abrupt, manifesting on day five. Mice exhibited similar degree of weight loss, but a varying degree of symptoms and clinical/histopathological scores. The number of mice used in this study was too small to determine whether this was a result of experimental variability or natural variability in outcomes. The variance in clinical and histopathological scores may be partially explained by viral titer, but there are likely other factors, such as the host immune response, that contribute to the variance observed. The observation of more severe disease in a subset of the K18-*hACE2* mice is distinct from other *hACE2*-expressing COVID-19 models, which typically observed only mild clinical symptoms (Bao *et al*., 2020)(Sun *et al*., 2020). This could be due to differences in the level of *hACE2 receptor* expression or tissue distribution. Based on these findings, the K18-hACE2 transgenic mice may be a particularly useful for studying the biological processes underlying the clinical symptoms of COVID-19 disease.

## Methods

### Challenge

Ten week-old Tg(K18-*hACE2*)2Prlmn (Jackson Laboratories) ((McCray *et al*., 2007) were challenged with 8 × 10^4^ TCID50 in 50 μl of Hong Kong/VM20001061/2020 (NR-52282, obtained from BEI Resources, NIAID, NIH) by intranasal route under ketamine/xylazine. Mock-infected animals received 50 μl DMEM. Mice were followed twice daily for clinical symptoms, which included weight loss, activity, fur appearance and posture, eye closure, and respiration. On day five all mice were euthanized. All mouse work was approved by the University’s Institutional Animal Care and Use, and all procedures were performed in the University’s certified animal Biosafety Level Three laboratory.

### Histology

Tissues were fixed in formaldehyde. Slides were scanned at 20X magnification. Histopathological scoring for lung tissue was done according to the guidelines of the American Thoracic Society (Matute-Bello *et al*., 2011).

### Viral titers

The left lobe of the lung was removed and placed in a disposable tissue grinder with 1 ml of serum free DMEM. Plaque assays were performed based on the protocol described in Baer and Hall, 2014. Briefly, Vero C1008, Clone E6 (ATCC CRL-1586) cells grown in DMEM (GIBCO 11995-040) supplemented with fetal bovine serum (FBS) were seeded into 12 well tissue culture plates at a concentration of 2 × 10^5^ cells/well the night before the assay. Lung homogenates were diluted in cold serum-free DMEM and serial dilutions added to the wells. The plate was incubated at 37°C, 5% CO_2_ for two hours to allow viral infection of the cells, shaking the plates every 15 minutes. After two hours, the media was replaced with a liquid overlay of DMEM, 2.5% FBS containing 1.2 % Avicel PH-101 (Sigma Aldrich) and incubated at 37°C, 5% CO_2_. After three days the overlay was removed, wells were fixed with 10% formaldehyde and stained with 0.1% crystal violet, and plaques enumerated to calculate viral PFU/ml.

### Bronchoalveolar lavage (BAL) fluid and flow cytometry

BAL was performed with 700 uL of PBS and cells were removed via centrifugation. Cells were stained then fixed in IC fixation buffer (eBioscience, 00-8222-49) and run and identified via Zombie NIR (Biolegend, 423105), CD45, Alexa Fluor 532 (eBioscience, 58-0459-42), CD11c, PE-Cy7 (Biolegend, 117317), CD11b, BV480 (BD, 566117), SIGLEC F, AF700 (eBioscience,56-1702-80), Ly-6c, FITC (Biolegend, 128005), Ly-6G, BV650 (Biolegend, 127641) on a Cytek Aurora Borealis at the University of Virginia flow cytometry core. Neutrophils are Zombie NIR-, CD45+, CD11C-, CD11B+, Ly-6G+, Eosinophils are Zombie NIR-, CD45+, CD11C-, CD11B+, Siglec F+, Inflammatory monocytes are Zombie NIR-, CD45+, CD11C-, CD11B+, Ly-6G-. Ly-6C ++. Data Analysis and figure generation was performed in Omiq and Graphpad Prism.

## Acknowledgements

The authors would like to gratefully acknowledge the advice and assistance of Angelina Angelucci and Young Hahn at the University of Virginia as well as Caitlin Woodson and Kylene Kehn-Hall at George Mason University. This work was supported by NIH grants R01 AI124214 to WAP, R01 AI146257 to SLB, and 5T32AI007496-23 to AND, and the University of Virginia’s Global Infectious Diseases Institute. JMS is a an iTHRIV Scholar, a program is supported in part by the National Center For Advancing Translational Sciences of the National Institutes of Health under Award Numbers UL1TR003015 and KL2TR003016. The content is solely the responsibility of the authors and does not necessarily represent the official views of the National Institutes of Health. The authors have no competing interests.

